# Development of the MAM model of schizophrenia in mice: Sex similarities and differences of hippocampal and prefrontal cortical function

**DOI:** 10.1101/295741

**Authors:** Kleanthi Chalkiadaki, Aggeliki Velli, Evangelos Kyriazidis, Vasiliky Stavroulaki, Vasilis Vouvoutsis, Ekaterini Chatzaki, Michalis Aivaliotis, Kyriaki Sidiropoulou

## Abstract

Schizophrenia is a debilitating disorder with complex and unclarified etiological factors. Sex differences have been observed in humans but animal models have only focused on male subjects. In this study, we report the establishment of the neurodevelopmental MAM model of schizophrenia in mice and compare the schizotypic-like characteristics and cognitive function in both sexes. Pregnant mice were injected with 26mg/kg(i.p.) of Methylazoxy-methanol acetate (MAM) or saline (5ml/kg) on gestational day (GD) 16 (MAM-16) or 17 (MAM-17). Behavioral, histological and electrophysiological and mass spectrometry-based comparative proteomic techniques were employed to assess the schizotypic-like characteristics and cognitive function of adult male and female offspring (MAM- or saline-treated). Female MAM-16, but not MAM-17 treated mice exhibited enhanced hyperlocomotion after acute administration of the NMDA receptor antagonist, MK-801, compared to saline treated mice. Male MAM-16, but not MAM-17 treated mice showed decreased pre-pulse inhibition of the acoustic startle reflex. Both male and female MAM-16 and MAM-17 treated mice exhibited reduced hippocampal (HPC) size and thinning of the prefrontal cortex (PFC), but only male MAM-16 treated mice showed decreased parvalbumin expression in HPC and PFC. Similarly, both male and female MAM-16 treated mice displayed impaired contextual fear memory, while only male MAM-16 treated mice exhibited deficits in the delayed alternation task. The neurophysiological mechanisms that underlie these cognitive functions were further investigated. Both male and female MAM-16 treated mice had significantly reduced long-term potentiation (LTP) in the HPC CA1 synapses, while only male MAM-16 treated mice exhibited decreased LTP in the PFC. Proteomic analyses of PFC lysates further showed significant MAM- and sex-dependent differences in regulation of protein expression. Our results demonstrate that while both male and female mice, prenatally exposed to MAM on GD16, display several core schizophrenia-like deficits and impairments in the hippocampus, only male MAM-treated mice have PFC-dependent cognitive deficits.

## Introduction

Schizophrenia (SZ) is a chronic, severe mental disorder that affects about 1% of the population (data from WHO 2018). Its symptoms are mainly categorized as positive (e.g. delusions, hallucinations), negative (e.g. avolition, blunted emotion) and cognitive (e.g. working memory deficits), and arise after late adolescence/early adulthood (Stahl 2013; Tandon et al. 2013). SZ is a multi-factorial disorder with several factors contributing to it, such as gene polymorphisms, epigenetic modifications and environmental factors (Gejman et al. 2010; Karayiorgou & Gogos 2014). However, the precise contribution and interactions of these factors remain unclear. The main hypotheses proposed for the pathophysiology of SZ are: a) the dopamine (DA) hypothesis, which supports that excessive DAergic transmission underlies the main symptoms of the disease (Davis et al. 1991) (for reviews see (Howes & Kapur 2009; Brisch 2014)), b) the glutamate hypothesis (Carlsson & Carlsson 1990), which suggests that NMDA receptor hypofunction in the brain is involved in the pathophysiology of SZ (Olney et al. 1999) (for reviews see (Nakazawa et al. 2012; Snyder & Gao 2013)) and c) the neurodevelopmental hypothesis (Marenco & Weinberger 2000; Lafargue & Brasic 2000; Fatemi & Folsom 2009; Owen et al. 2011) which proposes that SZ arises from insults that disturb central nervous system (CNS) development, leading to the observed pathologies in adulthood. Based on these hypotheses, animal models of SZ have been developed and have contributed significant information regarding the etiology and pathophysiology of the disease. A reliable animal model should present symptom homology (face validity), replicate the theoretical neurobiological base of the disease (construct validity) and show the expected pharmacological response or lack of it, after treatment with known antipsychotics or novel drugs tested (predictive validity).

The MAM model of SZ has been developed and characterized in rats. It utilizes the anti-mitotic agent methylazoxymethanol acetate (MAM) administered in pregnant rats at gestation day 17 (GD17) (Moore et al. 2006). It specifically targets neuroblast development without affecting glial cells (Cattaneo et al. 1995).This model is based on the neurodevelopmental hypothesis of SZ, thus satisfying the construct validity criterion (Jones et al. 2011). MAM-treated offspring show abnormalities resembling the schizotypic phenotype that arises after puberty. Specifically, MAM-treated rats exhibit enhanced locomotor activity in response to amphetamine (Moore et al. 2006), spontaneous hyperactivity to a novel environment (Pen et al. 2011) and impaired prepulse inhibition (PPI) of the acoustic startle reflex (Moore et al. 2006; Hazane et al. 2009; Pen et al. 2011), all considered equivalent to positive symptoms of SZ. Additionally, these animals show deficits in social behavior (Flagstad et al. 2004; Le Pen et al. 2006; Hazane et al. 2009) equivalent to the negative symptoms, as well as in cognitive tasks, including impairments in spatial learning and memory (Gourevitch et al. 2004; Hazane et al. 2009; Ratajczak et al. 2015; Gastambide et al. 2015; Gill et al. 2017), cognitive flexibility (Moore et al. 2006) and problem-solving (Featherstone et al. 2007). MAM-17 rats also have reduced thickness in cortical and subcortical regions (Moore et al. 2006; Flagstad et al. 2004; Featherstone et al. 2007; Matricon et al. 2010; Hradetzky et al. 2012), dysregulated dopamine (DA) system (Grace 2017) and parvalbumin (PV) loss in ventral subicullum, HPC and PFC (Penschuck et al. 2006; Lodge et al. 2009; Gill & Grace 2014). All of the above data support the face validity of the model.

On the other hand, genetic models, almost exclusively developed in mice, have significantly contributed to understanding the genetic endophenotypes of SZ, but cannot replicate the complexity of the disorder (Birnbaum & Weinberger 2017).

Given the etiological complexity of the disease, the development of more advanced animal models of SZ is essential. Genetic and environmental factors need to be integrated by developing animal models based on gene-environment interaction (Papaleo et al. 2012; McOmish et al. 2014; Ayhan et al. 2009).

Furthermore, several sex differences with regards to symptoms manifestations have been observed in humans, including earlier onset of the disease for males, differential symptoms and response to treatment (Leger & Neill 2016). However, very few studies using animal models have focused on females. Thus, the possible mechanisms that underlie the sex differences remain unknown.

Our study reports the development of the MAM model in both male and female mice revealing both sex similarities and differences. In particular, we have noted that injection of MAM on GD16 better reproduces the positive symptoms and the key histological alterations in both male and female mice, compared to GD17 MAM injection employed in rats. In addition, we find that HPC function is significantly deteriorated in both male and female MAM-treated mice. However, female MAM-treated mice exhibit decreased vulnerability of PFC function deficits, as shown with behavioral, electrophysiological and proteomic analyses.

## Methods and Materials

### Animals and MAM treatment

All experiments were conducted in adult (>3 months old) C57BL/6 male and female offspring of pregnant dams treated with either saline or MAM. Mice were housed in groups (3-4 per cage) and provided with standard mouse chow and water ad libitum, under a 12 h light/dark cycle (light on at 7:00 am) with controlled temperature (23 +/- 1 Celsius). All procedures were performed according to protocols approved by the Research Ethics Committee of the University of Crete and obey the European Union ethical standards outlined in the Council Directive 2010/63EU of the European Parliament on the protection of animals used for scientific purposes.

Time pregnant dams received intraperitoneal (i.p.) injections of MAM (MRI global, Kansas City, MO) (26mg/kg) or saline (1ml/kg) on GD 16 or 17. According to species comparison of Carnegie stages of embryonic development (https://embryology.med.unsw.edu.au/embryology/index.php/Carnegie_Stage_Comparison), GD16 rather than GD17 of mice corresponds better to the GD17 of rats, regarding the morphological development of the embryo. In addition, the latest predictive model (Workman et al. 2013) proposes that equivalent maturation stages of brain development between mouse and rat differ 1-2 days, with the latter maturing later. Based on these data, we initially tested mice treated on both GDs in validation experiments, referred from now on as MAM-16 and MAM-17 mice. We continued our study using the offspring with the most intense phenotype to reduce the number of animals used. All injections were conducted between 01.00-03.00p.m. Time pregnant BALB/c dams were used as foster mothers, until pups were weaned on day 25. All tests were performed in adulthood; that is 90-150 days old.

### Behavioral experiments

All behavioral experiments were conducted between 10.00a.m.-5.00p.m.

### MK-801 challenge

Locomotor hyperactivity in response to MK-801 was assessed in a total of 30 female animals, 10 mice per group (saline, MAM-16, MAM-17). Animals were moved to the experimentation room, 1 h prior the test, for acclimation. The experiment took place under low illumination conditions. Animals were placed in an open field arena (45×45×45cm) for a 3-hour habituation period. After subcutaneous (s.c.) saline injection (1ml/kg), they were placed back to the arena for 30 minutes, and finally they received an i.p. injection of MK-801 and placed back for 90 more minutes. Locomotor activity was video-recorded during the whole task, using a recording camera. All videos were analyzed with ANY-maze tracking software. For each animal, total distance traveled was measured throughout the experiment in 10min bins.

### Prepulse Inhibition (PPI) of the acoustic startle reflex

The expression of PPI of the acoustic startle reflex was assessed in a total of 27 male mice (10 saline,12 MAM-16 and 7 MAM-17). We used a custom-made PPI apparatus, which consisted of a mini-chamber (3×7×2cm), located inside a ventilated plywood sound attenuating box, dimly lit. Animals were habituated to the mini-chamber prior to testing for 3 non-consecutive days, 10 minutes each day, to reduce anxiety levels. All acoustic noise bursts and background noise were delivered through a computer, connected to a speaker located inside the box. Throughout the habituation sessions a background noise level of 68dB was maintained. At testing day, after a 7-minute acclimation period (68dB background noise), animals were submitted to a series of 5 startle stimuli, referred as pulse-alone (115dB, duration 50ms), with inter-trial intervals of 20-25 seconds, aiming to accustom the animals to the startle pulses. Subsequently, animals received 15 pulse-alone (115dB, 50ms), 15 prepulse+pulse, where the prepulse was 6-12 dB above background, had duration of 20ms and inter-stimulus interval between the prepulse and the pulse 40ms, and 10 prepulse-alone stimuli, pseudo-randomly presented every 20-25 sec. The behavior of the animal was monitored throughout the task, using a camera placed inside the box. Animal response to each noise burst was tracked with OpenVision control tracking software, exported as image processing data and analyzed with custom-made Matlab code (http://github.com/NBLab/PPI). In order to calculate the startle response, 5 image frames before each sound (1 sec) and 1 image frame after each sound (20msec) were analyzed for animal movement. Animal movement was counted as the number of pixels that changed from one frame to the other. Startle amplitude was calculated as the ratio of the number of moved pixels after the sound to the number of moved pixels before the sound. Mean amplitude of startle response to pulse alone (P) and Prepulse+pulse (PP+P) trials, was calculated for each animal. The level of PPI was assessed by expressing the Prepulse+pulse response amplitude as a percentage decrease from pulse alone response amplitude, using the following formula: %PPI=100-[100 × (PP/P)].

### Contextual fear conditioning

The contextual fear conditioning paradigm was used to assess fear memory formation, as outlined before (Nikoletopoulou et al. 2017). Specifically, animals were transferred to the experimentation area 1 h prior the experiment for acclimation and at every single day of the task, both groups of animals (saline and MAM-treated) were used. On the 1st day of the experiment (training day), mice were placed in the fear-conditioning chamber (MedAssociates, St Albans, VT, USA), which was controlled through a custom-made interface connected to the computer, for 10 minutes. After a 7 min of habituation to the chamber, a mild electrical foot shock (0.7mA, 1sec) was delivered and the animal remained for another 3 minutes. The following day, mice were returned to the training chamber but did not receive any electrical shock. The activity of each animal was recorded and freezing behavior was analyzed with the JWatcher software (http://www.jwatcher.ucla.edu/). Every 5 seconds the observer scored the behavior of the animal (moving or not moving-freezing) and the freezing behavior was calculated using the formula: %freezing=[not moving/(moving + not moving)].

### Delayed alternation task in the T-maze

The T-maze apparatus includes a start arm and two goal arms (45×5cm each). The delayed alternation task in the T-maze is a classic behavioral task used for the study of working memory (Konstantoudaki et al. 2018). Mice were initially handled by the experimenter for about a week, food-restricted so that animals maintained 85-90% of their initial weight and then habituated in the T-maze apparatus for 2 days. Mice were subjected to 10-trial sessions, 3 sessions per day. At the first trial of each session, mice were allowed to freely choose between the right or left goal arms. In the following trials, mice had to alternate the goal arms in order to receive reward, initially with no temporal delay between the trials. Once they reached a pre-defined criterion for the alternation procedure (i.e., 2 consecutive sessions of ≥70% correct choices (performance), delays were introduced starting at 5 seconds and increasing by 5 seconds when the criterion for each delay was achieved, until mice completed the alternation procedure with a 15 seconds delay. Upon successful completion of the above mentioned trials, mice were tested for 2 days, 3 sessions a day, in a random set of delays ranging from 5 seconds to 25 seconds in order to better test performance in the working memory task.

### Tissue preparation

For the histological experiments, mice were deeply anesthetized with 20mg/ml avertin (250mg/kg, i.p.), perfused with phosphate buffer saline (PBS) and subsequently with ice cold phosphate-buffered 4% paraformaldehyde. Brains were removed, post-fixed for 24 hours and preserved in PBS-azide at 4°C, until slicing with a vibratome (VT1000S, Leica Microsystems, Wetzlar, Germany). 40-µm-thick coronal slices were cut sequentially in sets of four sections. 3-5 sections per animal were used for neuroanatomy and immunohistochemistry experiments, which corresponded to different rostro-caudal levels of the brain.

### Neuroanatomy based on Nissl staining

Slices containing the PFC, HPC or barrel cortex (BC) were stained with cresyl violet, as previously described (Konstantoudaki et al. 2016). Briefly, sections were incubated in xylene (5min), 90% and 70% ethanol solutions (3min), dH_2_O, followed by 10-min incubation in 0.1% Cresyl Violet solution. Sections were then dehydrated with increasing concentrations of ethanol (70%, 90%,100%), incubated in xylene for 5min and coverslipped with permount. Sections containing PFC were taken from Bregma 2.22 to 1.70 mm, sections containing dorsal HPC were taken from Bregma - 1.34 to −2.06mm and finally, sections containing BC corresponded to three different levels: Bregma -0.94 to -1.22mm, -1.34 to -1.70mm and -1.82 to −2.06mm. This categorization aimed to minimize the bias in the measurements, if BC thickness changed along the rostro-caudal axis. Images from whole sections were obtained in 5x magnification of a light microscope (Axioskop2FS, Carl Zeiss AG, Oberkochen, Germany). Multiple overlapped pictures were taken for each slice and merged using Adobe Photoshop CS6 software. We measured cortical thickness for PFC and BC as well as the horizontal and vertical dimensions of HPC. According to the literature, three subdivisions of PFC have been recognized, in terms of cyto-architecture, chemo-anatomy, connectivity and function (Heidbreder & Groenewegen 2003; Etkin et al. 2011; Giustino & Maren 2015), namely the anterior cingulate (ACC), Prelimbic (PrL) and Infralimbic (IL) cortices. The width of each different subdivision was measured from midline to the beginning of the white matter. For HPC, a horizontal straight line was taken from the dorsal tip of the third ventricle, reaching the corpus callosum, for the horizontal plane measurement and a vertical straight line was taken in the middle of the natural curve the structure forms, extending until the beginning of thalamus. The exact position of the BC was identified using the online mouse brain atlas (http://www.brain-map.org/). Two vertical lines from the edge of corpus callosum, reaching layer I of the cortex (650µm in between distance) were used for thickness measurement. To ensure the accuracy of the measurements, the two lines were taken 260µm inwards the approximate borders of BC. The average of the two measurements was calculated and used for the analysis. All area measurements were conducted manually, using Adobe Photoshop CS6 software.

### Fluorescent immunohistochemistry

Free floating sections, adjacent to the Nissl-stained sections, containing PFC or HPC (as described above) were stained using indirect fluorescent immunohistochemistry for detection of PV–containing interneurons. Briefly, sections were rinsed with Tris-buffered saline (TBS, 1M), blocked for 90 minutes with 10% fetal bovine serum (FBS), 0.4% Triton in TBS-Tween 0.01% and incubated with primary antibody (rabbit-polyclonal anti-PV, 1:3000, PV27, Swant, Inc., Switzerland) in 5% FBS, 0.2% Triton in TBS-Tween 0.01%, overnight at 4°C. Sections werethen incubated in secondary antibody (Goat-anti-rabbit, Alexa-488 conjugated, 1:500, Thermo Fisher Scientific, Inc., U.S.A.) for 2 hours at room temperature. They were subsequently rinsed with TBS-Tween 0.01% and incubated with Propidium Iodide in TBS for 7 minutes, after a 30 minute incubation with RNAseA (Quiagen, Inc., U.S.A.) in TBS. Finally, sections were rinsed with TBS, mounted onto slides and coverslipped with Mowiol solution. After PV immunostaining, images were obtained with a Confocal microscope (Leica TCS, SP1, Leica Microsystems, Mannheim, Germany) using the 10x objective. Multiple overlapped pictures were taken for each region of interest and merged using Adobe Photoshop CS6 software. For HPC containing sections, PV-positive cells were counted manually for the CA1 area. For PFC-containing sections, PV-positive cells were summed in the ACC and PrL areas. Merged images were cropped in the regions of interest, according to mouse brain atlas and the background color of each cropped image was converted to black, while the cells were colored green. Images were then loaded into Matlab, where the number of green cells was counted in each image, regardless of staining intensity.

### Electrophysiology

Electrophysiological experiments were performed using the *in vitro* slice preparation as previously described (Konstantoudaki et al. 2016). Mice were decapitated under halothane anesthesia. The brain was removed immediately and placed in cold, oxygenated (95% O_2_ / 5% CO_2_) artificial cerebrospinal fluid (aCSF) containing (in mM): 125 NaCl, 3.5 KCl, 26 NaHCO_3_, 1 MgCl_2_ and 10 glucose (pH= 7.4, 315 mOsm/l). The brain was blocked and glued onto the stage of a vibratome (Leica, VT1000S, Leica Biosystems GmbH, Wetzlar, Germany). 400-µm-thick brain slices containing either the PFC or the HPC were taken and were transferred to a submerged chamber, which was continuously superfused with oxygenated (95% O_2_ / 5% CO_2_) aCSF containing (in mM): 125 NaCl, 3.5 KCl, 26 NaHCO_3_, 2CaCl_2_, 1 MgCl_2_ and 10 glucose (pH= 7.4, 315 mOsm/l) at room temperature. After 1-2 hours of equilibration, slices were transferred to a submerged recording chamber, which was continuously superfused with oxygenated (95% O_2_ / 5% CO_2_) aCSF(same as constitution as the one used for maintenance of brain slices), at room temperature. Extracellular recordings were conducted in one or two slices per animal for each region studied.

For HPC recordings, extracellular recording electrodes filled with NaCl (2M) were placed on pyramidal layer of CA1 subregion of dHPC containing slices (Bregma-1.34 to −2.06mm). Platinum/iridium metal microelectrodes (Harvard apparatus UK, Cambridge, UK) were placed on CA1 subregion, about 300µm away from the recording electrode and near CA3 subregion. For PFC recordings, both the recording and stimulating electrodes were placed on layer II/III, (in-between distance 300µm), of PFC-containing slices (Bregma 2.22 to 1.70 mm). Stimulation of the regions evoked field excitatory postsynaptic potentials (fEPSPs) that were amplified using a Dagan BVC-700A amplifier (Dagan Corporation, Minneapolis, MN, USA), digitized using the ITC-18 board (Instrutech, Inc, Longmont, CO, USA) on a PC, using custom-made procedures in IgorPro (Wavemetrics, Inc, Lake Oswego, OR, USA). The electrical stimulus consisted of a single square waveform of 100µsec duration given at intensities of 0.05-0.3mA, generated by a stimulator equipped with a stimulus isolation unit (World Precision Instruments, Inc, Sarasota, FL, USA).

### Data acquisition and analysis

Data were acquired and analyzed using custom-written procedures in IgorPro software. The fEPSP amplitude was measured from the minimum value of the synaptic response (4-5 ms following stimulation) compared to the baseline value prior to stimulation. Both parameters were monitored in real-time in every experiment. A stimulus-response curve was then determined using stimulation intensities between 0.05 and 0.3mA. For each different intensity level, two traces were acquired and averaged. Baseline stimulation parameters were selected to evoke a response of 1mV. For the long-term potentiation experiments, baseline responses were acquired for at least 20 minutes. Then two theta-burst stimulation trains (5 bursts of 100Hz/5Hz) were applied in dHPC-containing slices, while three 1second tetanic stimuli (100Hz) with an inter-stimulus interval of 20 seconds were applied in PFC containing slices. Finally, responses were acquired for at least 30 minutes and 50 minutes post theta-burst and post-tetanus, for HPC and PFC recordings, respectively. Synaptic responses were normalized to the average 10 minutes pre-theta burst/pre-tetanic fEPSP.

### Bottom-up comparative proteomic analysis

Fresh PFC tissues of saline and MAM-treated mice of both sexes were frozen (−80 °C) and homogenized in lysis buffer (50mM HEPES, 150mM NaCl, 1% Glycerol, 1% Triton, 1,5 mM MgCl2, 5mM EGTA, 1:1000 inhibitor cocktail). Protein extracts were fractionated by SDS-PAGE (40 µg of sample protein in each lane, 10% acrylamide separating gel, 4% acrylamide stacking gel, electrophoresis at 140 V for 80 min) and stained with the MS-compatible blue silver staining protocol (Giovanni et al. 2004). Each lane corresponding to MAM-16 and saline animals was excised completely from the gel into multiple gel slices and in-gel trypsin digested separately (Shevchenko et al. 1996). The resulted peptide mixture was dried in a speed-vacuum centrifuge and re-constituted in 0.5% formic acid aqueous solution. Protein identification and relative quantitation by nLC-ESI-MS/MS was done on a LTQ-Orbitrap XL coupled to an Easy nLC (Thermo Fisher Scientific, Inc., U.S.A.). The sample preparation and the LC separation were performed as described previously (Aivaliotis et al. 2007), with minor modifications. Briefly, the dried peptides were dissolved in 0.5% formic acid aqueous solution, and the tryptic peptide mixtures were separated on a reversed-phase column (ReprosilPur C18 AQ, Dr. Maisch), fused silica emitters 100 mm long with a 75 µm internal diameter (Thermo Fisher Scientific, Inc., U.S.A.) packed in-house using a pressurized packing bomb (Loader kit SP035, Proxeon). Tryptic peptides were separated and eluted in a linear water-acetonitrile gradient and injected into the MS as described in earlier work (Aivaliotis et al. 2007; Aivaliotis et al. 2009). The nLC-MS/MS raw data were loaded in Proteome Discoverer 1.3.0.339 (Thermo Fisher Scientific, Inc., U.S.A.) and run using Mascot 2.3.02 (Matrix Science, London, UK) search algorithm against the Mouse theoretical proteome (Apweiler 2009). For protein identification, the following search parameters were used: precursor error tolerance 10 ppm, fragment ion tolerance 0.8 Da, trypsin full specificity, maximum number of missed cleavages 3, methionine oxidation as variable modifications. Relative protein quantitation was performed using Scaffold_4.4.8 (Proteome Software Inc.) based on the MS intensities of the identified peptides.

### Statistical Analyses

Results were analyzed for statistical significance using the SPSS software. All graphs report the average ± standard error of the mean. One-way or multi-way ANOVA were performed according to the study. The Kruskal-Wallis or Mann-Whitney tests were used when data did not fit a normal distribution. Statistical significance was set at p<0.05 and is indicated with an asterisk (*) in the graphs. Trend for statistical significance was set at p<0.1 and is indicated with (#) in the graphs.

## Results

The MAM model has not been previously studied in mice. Therefore, we first conducted validation experiments, in three groups of animals: a) mice that were in injected with saline at GD16 (saline group), b) mice that were injected with MAM at GD17 (MAM-17 group), which is the same injection time-point as in rats, and c) mice that were injected with MAM at GD16 (MAM-16 group), the time point that we predict will have a stronger phenotype and similar to the rat MAM model. The validation experiments included the investigation of histological alterations of the HPC, PFC and BC, behavioral tests of ‘schizotypic-like’ symptoms (MK-801-induced hyperlocomotion in female mice and PPI of the acoustic startle reflex test in male mice), due to sexual dimorphic results reported in the literature (see discussion below), as well as immunostaining for the PV protein, in both sexes.

### Morphological alterations in brain regions after MAM exposure

Histological analysis, using Nissl staining, revealed anatomical alterations in cortical and subcortical brain regions in MAM treated mice. For HPC, a decrease in the vertical plane of the hippocampal formation in female MAM-17 mice (Kruskal-Wallis test, *p*=0.03) and a trend towards a decrease in female MAM-16 mice (Kruskal-Wallis test, *p*=0.08) were observed (Figure 1A). Similar effects were identified in male mice, in which both MAM-16 and MAM-17 mice exhibit significantly reduced vertical dimension, compared to saline-treated mice (Kruskal-Wallis test, *p*=0.01 and p=0.03, respectively). At the horizontal plane, analysis revealed a significant effect of MAM treatment in female mice for both MAM-16 (Kruskal-Wallis test, *p=0.006*) and MAM-17 group (Kruskal-Wallis, *p*=0.02) but no significant alterations were observed in either group of MAM-treated male mice, compared to saline-treated mice (Kruskal-Wallis test, *p*=0.70 and *p*=0.30, for MAM-17 and MAM-16 respectively) (Figure 1A).

**Figure 1:**
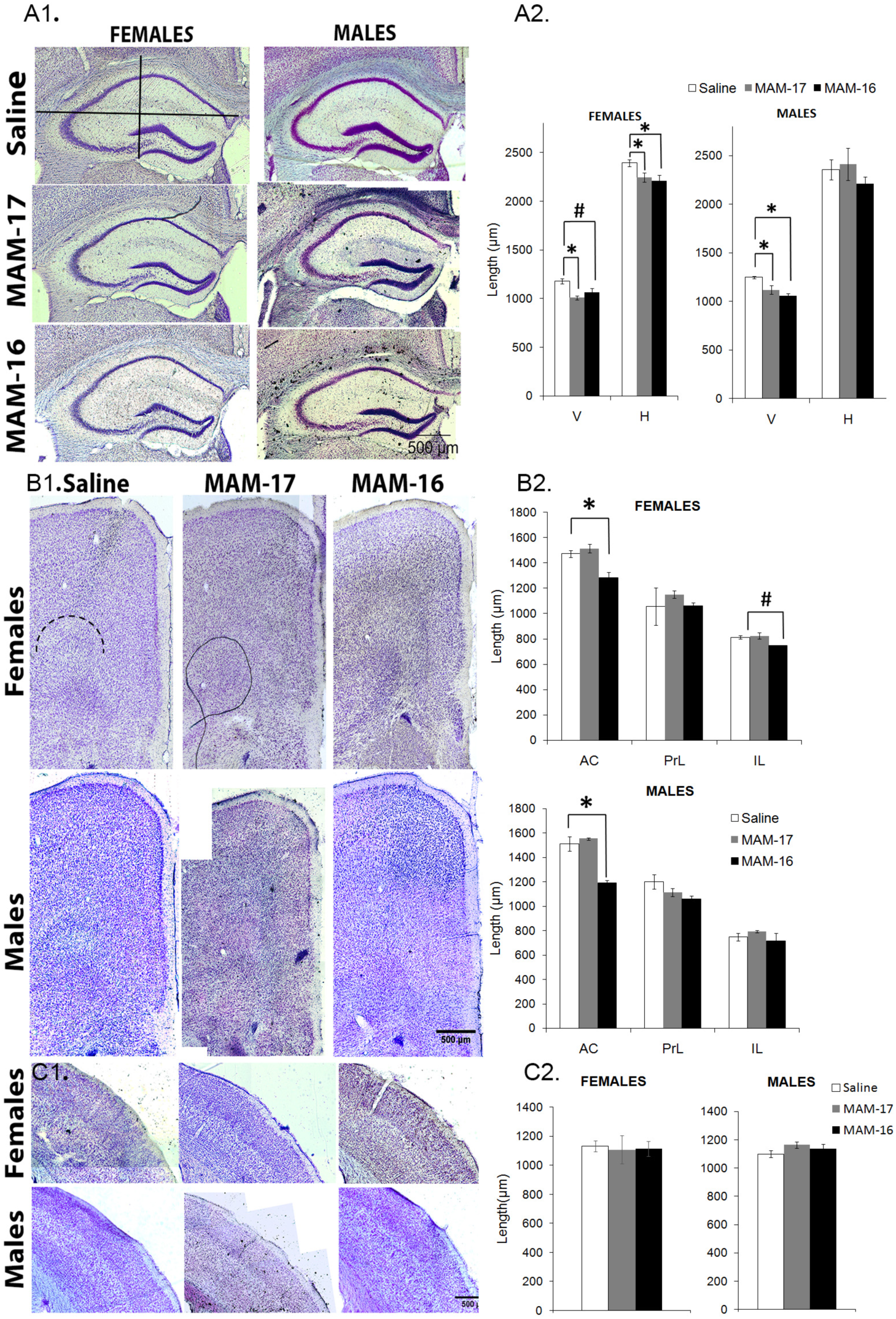
Anatomical alterations in cortical and subcortical brain regions. Representative images of coronal sections containing the dorsal hippocampus of Saline- and MAM-exposed male and female mice. Bar graphs showing the vertical (V) and horizontal (H) plane measurements in each group. A significant effect of MAM treatment on the vertical plane was observed in both female and male MAM-17 mice (Kruskal-Wallis test, p=0.05 and p=0.016, respectively), in MAM-16 male mice (Kruskal-Wallis test, p=0.014) and a trend for decrease in MAM-16 female mice (Kruskal-Wallis test, p=0.08). A significant decrease in the horizontal dimension was only observed in MAM-16 and MAM-17 female mice (Kruskal-Wallis test, p=0.006and p=0.02, respectively). No alterations in MAM-exposed male mice (Kruskal-Wallis test, p=0.72 and p=0.31, MAM-17 and MAM-16, respectively). Representative images of coronal sections containing the prefrontal cortex. Bar graphs showing the width of each subregion. A significant thinning of ACC was observed in MAM-16 female and male mice (Kruskal-Wallis test, p = .02 and p = .006, respectively), no alterations in PrL cortex thickness, (Kruskal-Wallis test p = .2 for both sexes and groups, compared to control animals). A trend for thinning of IL cortex only in MAM-16 female mice (Kruskal-Wallis test, p =.06). Representative images of coronal sections containing the Barrel cortex. F) Bar graphs showing the absence of differences in barrel cortex thickness between saline and MAM-exposed mice, in both sexes. MAM: methylazoxymethanol acetate, ACC: Anterior cingulate cortex, PrL=Prelimbic and IL=Infralimbic). [males: n(saline) =4, n(MAM-17)=3, n(MAM-16)= 5, females: n(saline) =6, n(MAM-17)=3, n(MAM-16)=5]

For the PFC, we analyzed the width of ACC, PrL and IL. ACC width was found statistically decreased in MAM-16 female and in MAM-16 male mice, compared to their respective saline-treated mice (Kruskal-Wallis test, *p*=0.02 and *p*=0.006, respectively), whereas no alterations were observed in MAM-17 female and male mice (Kruskal-Wallis test, *p*=0.43) (Figure1B). No significant alterations were found in the width of PrL subregion between MAM-treated and saline-treated animals (Kruskal-Wallis test, *p*=0.20 for both females and males [MAM-16/MAM-17]). Additionally, the analysis of the IL cortex width showed a trend for thinning only in female MAM-16 mice (Kruskal-Wallis test, *p*=0.06) and no alterations in male MAM-16 mice (Kruskal-Wallis test, *p*=0.35) or MAM-17 mice groups (Kruskal-Wallis test, *p*=0.71 and *p*=0.60, for females and males,respectively).Finally, no significant differences were identified between MAM-16 nor MAM-17 and saline-treated mice in BC of females (Kruskal-Wallis test, *p*=0.80 and 0.60, MAM-16 and MAM-17, respectively) and or males (Kruskal-Wallis test, *p*=0.60 and.1 for MAM-16 and MAM-17, respectively) (Figure 1C).

### Schizotypic-like symptoms in MAM mice

We subsequently examined the possible schizotypic-like symptoms of the MAM-treated mice at the behavioral level, such as the enhanced locomotor activity in response to an MK-801 challenge and the reduced PPI of the acoustic startle reflex. Animals were separated by sex, in an effort to reduce the number of subjects used. In particular, female mice were examined for their locomotor activity in response to MK-801, since females are more sensitive in this task (Andiné et al. 1999). Male mice were used in the PPI task, as there have been indications for control females exhibiting reduced levels of PPI (Kumari et al. 2004; Matsuo et al. 2016).

In female mice, no differences were observed among the groups with regards to locomotor activity during the first three hours of the test or the 30min after saline (s.c.) injection. In all groups, locomotor activity decreased over time, indicative of normal habituation in the apparatus. Administration of MK-801 (0.2 mg/kg, i.p.) resulted in enhanced locomotor activity in all groups (repeated measures ANOVA,*F*_(2,27)_=3.47, *p*=0.04). Post-hoc analysis showed a significant effect of MAM treatment in the MAM-16 group (*p*=0.04), but no significant effect of MAM exposure in the MAM-17 group (*p*=0.5) (Figure 2A). In addition, analysis of the distance travelled, showed that MAM-16 mice had already travelled significantly longer inside the apparatus 50 minutes after the MK-801 injection (post-hoc LSD test, *p*=0.02), compared to saline-treated mice, while MAM-17 mice did not seem to differ significantly from the control group (post hoc, *p*=0.02) (Figure 2A).

**Figure 2:**
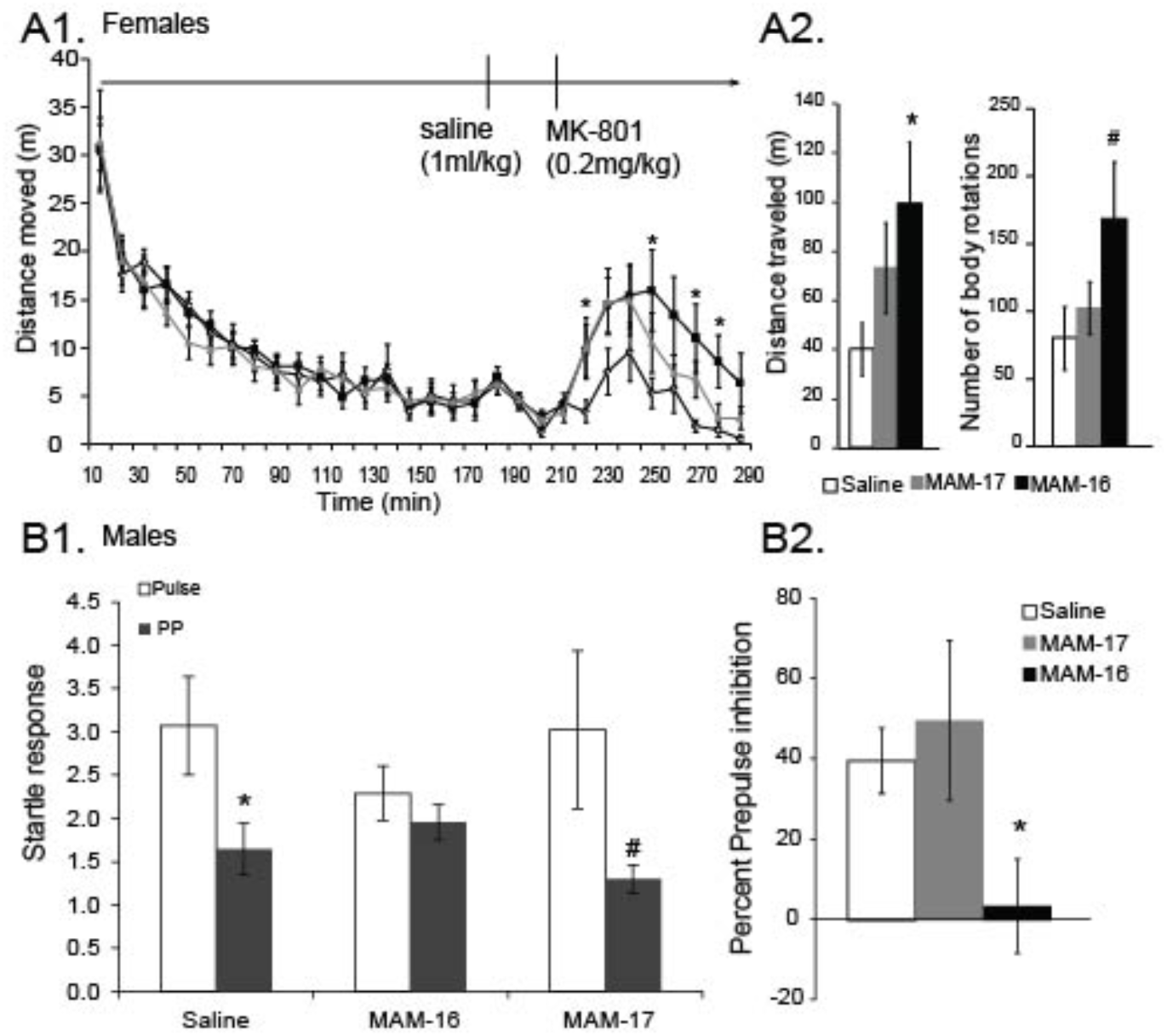
Schizotypic-like’ phenotype of male and female MAM-treated mice. Locomotor hyper-activity of female mice after systemic treatment with MK-801. (Top) Line plot showing the locomotor activity in the open field, expressed as distance (m) travelled, during habituation phase (0-180min), after saline injection (180min-210min) and after treatment with 0.2mg/kg of MK-801 (210min-300min). Post hoc analysis revealed a significant effect of MAM treatment in MAM-16 (p=.04), but not in MAM-17 mice (p=.5). MAM-16 mice exhibited significantly increased hyper-locomotion at 40, 70, 90 and 100 minutes after the acute MK-801 injection, compared to saline mice (repeated measures ANOVA p<.01 at all time points, n(Saline)=10, n(MAM-16)=10, n(MAM-17)=10). (Bottom) Bar graph showing the mean total distance travelled for each group, during the first 50 minutes after the MK801 injection. MAM-16mice moved significantly longer that saline mice (p = .15). Prepulse inhibition of the acoustic startle reflex of male mice. (Top) Bar graphs showing the mean startle response of each group at pulse alone (Pulse) and prepulse – pulse (PP) trials. Responses in different prepulse-intensity trials (72dB and 80dB) were averaged. No significant differences were observed in startle response between the groups (Kruskal-Wallis test, p = .43). (Bottom)The percent of PPI was significantly decreased in MAM-16 mice (Kruskal-Wallis test, p= .035), but not in MAM-17 mice. [n(Saline)=8, n(MAM-16)=10, n(MAM-17)=7]

In male mice, no significant differences were found in the startle response among the three groups (Kruskal-Wallis test, *p*=0.43) (Figure 2B). Saline-treated mice significantly inhibited their startle reflex in response to the prepulse-pulse tone (*p*=0.01), and so did the MAM-17 mice (*p*=0.06). However, MAM-16 group did not inhibit the startle reflex in response to the prepulse-pulse tone (*p*=0.20). Instead, there was a significant increase in the startle response (Kruskal-Wallis test, *p* =0.40). When all three groups were tested for their PPI index, we found a significant decrease in MAM-16 group, compared to saline (Kruskal-Wallis test, *p*=0.03), but normal expression of PPI in MAM-17 mice (Kruskal-Wallis test, *p*=0.14) (Figure2B).

### PV expression in MAM-16 mice

Based on our behavioral and anatomical results that showed a stronger ‘schizotypic’ phenotype of MAM-16, compared to MAM-17 mice, we continued our experiments with MAM-16 mice, because they had a stronger phenotype compared to MAM-17 mice. Decreased PV expression has been identified as a histological marker of SZ from post-mortem studies (Lewis et al. 2001; Zhang & Reynolds 2002; Wang et al. 2011). Using fluorescent immunohistochemistry for the PV protein, we counted the number of PV-positive cells in brain slices containing the HPC or PFC. Our analysis, revealed a sex-selective decrease in PV-positive interneurons in the brain of MAM-16 mice. In the CA1 HPC region, the analysis showed no alterations in the number of PV-positive cells in female MAM-16 mice (Mann-Whitney test*, p*=0.52), while in male MAM-16 mice a statistically significant decrease was observed in the number of PV-positive cells (Mann-Whitney test, *p*=0.02) (Figure 3A). Similarly, no alteration in the number of PV-positive cells of PFC was observed in MAM-16 female mice (Mann-Whitney test, *p*=0.50), but decreased number of PV-positive cells in MAM-16 male mice (Mann-Whitney test, *p*=0.05) (Figure 3A).

**Figure 3:**
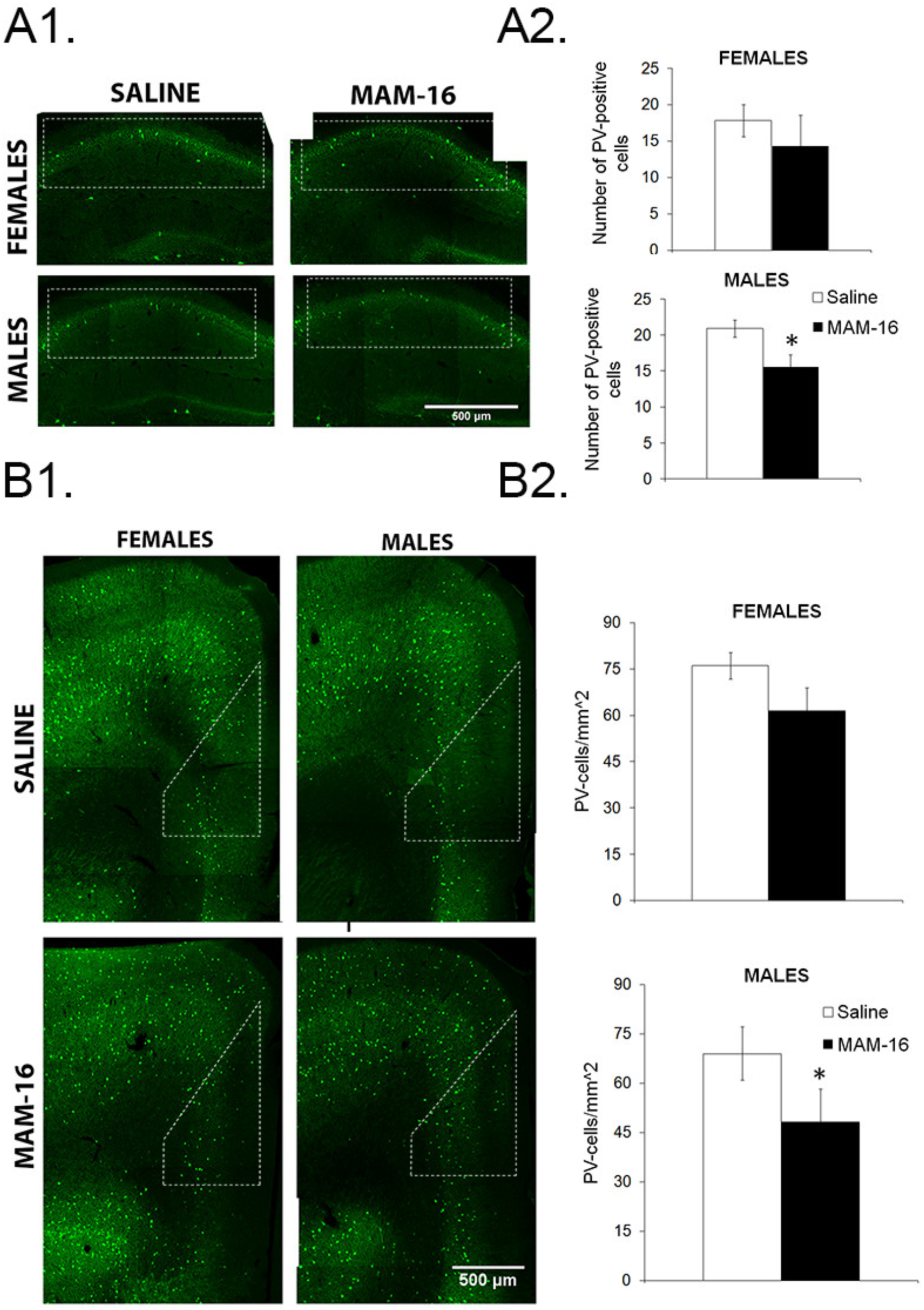
Parvalbumin expression in HPC and PFC of MAM-16 mice. A) (Left) Representative images of coronal hippocampal sections from saline and MAM-16 mice. White frames indicate the borders of CA1 region. (Right) Bar graphs showing a significant reduction in the number of PV-positive cells in CA1 of male MAM-16 mice (Mann-Whitney test, p= .04), but not of female MAM-16 mice (Mann-Whitney test, p= .2) B) (Left) Representative images of coronal sections containing PFC, from saline and MAM-16 mice. (Right) Bar graphs showing a significant reduction in the number of PV-positive cells in PFC of male MAM-16 mice (Mann-Whitney test, p= .04), but not in female MAM-16 mice. [males: n(saline)=3, n(MAM-16)=4, females: n(saline)= 4, n(MAM-16)=5].

### Both male and female MAM-16 mice exhibit HPC function deficits

The MAM model in rats has also been shown to affect HPC function (Penschuck et al. 2006; Moore et al. 2006; Matricon et al. 2010; Hradetzky et al. 2012; Gill & Grace 2014; Sanderson et al. 2012; Snyder et al. 2013). To assess HPC function in our model, the contextual-fear conditioning paradigm was used in both female and male mice. Mice were trained and tested for fear memory 24 hours later. Both female and male MAM-16 mice showed a statistically significant decrease of their freezing behavior, compared to saline mice (*t*-test, *p* =0.03 and *p*=0.02, females and males, respectively) (Figure 4A-Β), suggesting that prenatal MAM exposure can cause deficits in contextual fear memory of both male and female mice. We next investigated synaptic transmission and plasticity in hippocampal CA3 to CA1 synapses. We found that the fEPSP was similar for both females and males in response to increasing current stimulation (repeated measures ANOVA, F_(1,14)_=0.7, p=0.2 for females, F_(1,14)_= 0.5, p=.4 for males) (Figure 4C-D). Theta-burst stimulation induced LTP in both male and female saline-treated mice. MAM-16-treated mice, both males and females, exhibited significant reduction in the LTP (repeated measures ANOVA, F_(1,14)_ =10.1, p=0.001 for females and F_(1,14)_= 8.2, p=0.001 for males) (Figure 4E-F).

**Figure 4:**
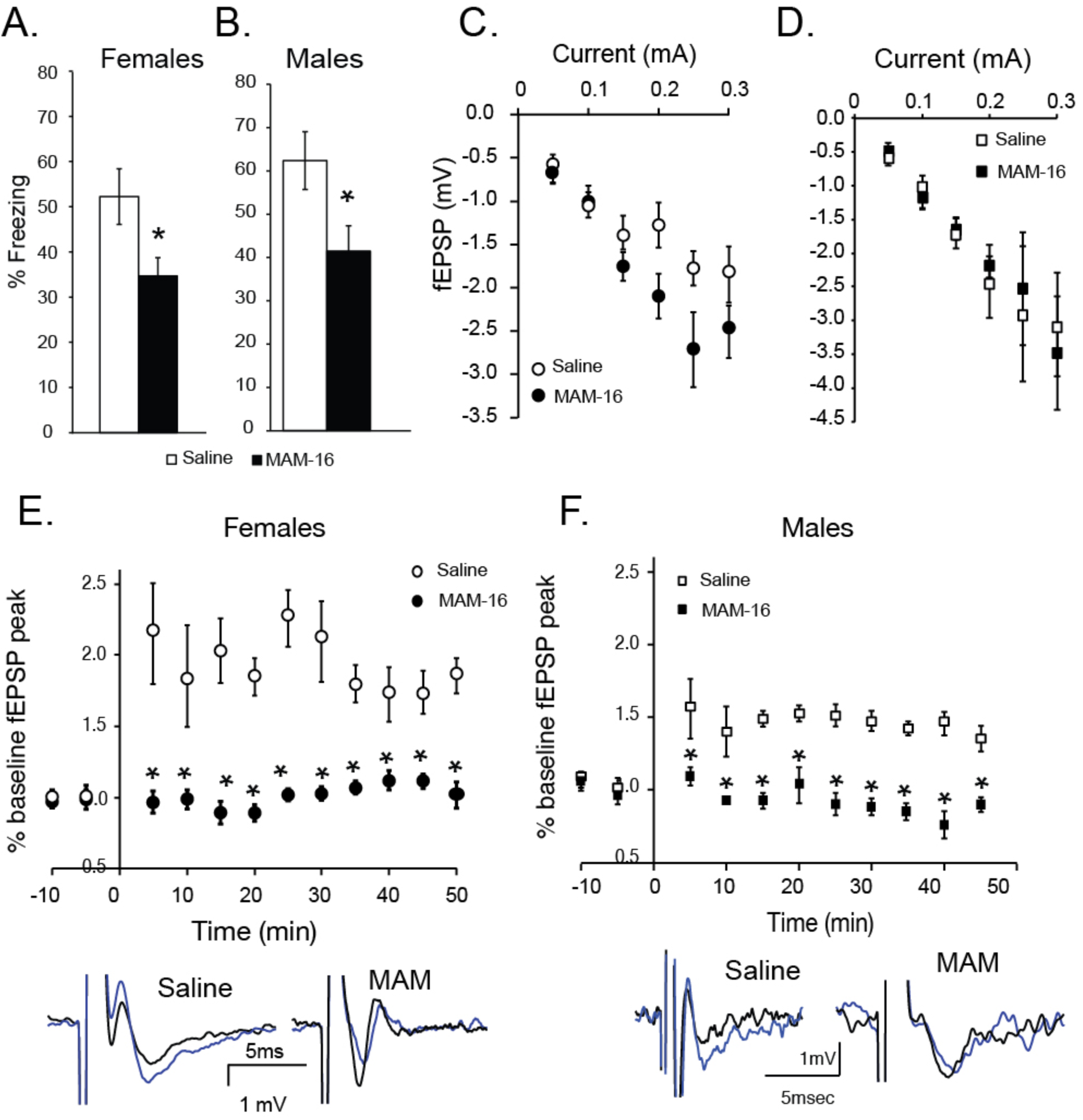
HPC function in male and female MAM-16 mice. A-B) Graphs showing reduced freezing behavior 24hours following training in the contextual fear conditioning task (t-test, p=0.03 and 0.02, males and females respectively) [males: n(Saline)=8, n(MAM-16)=11, females: n(Saline)=8, n(MAM-16)=9]. C-D) Graphs showing the fEPSP in response to increasing current stimulation. No significant differences are observed between saline and MAM-treated mice in either females (repeated measures ANOVA,F_(1,14)_=0.7, p=.2)or males(repeated measures ANOVA, F_(1,14)_= 0.5, p=.4 for males) [males: n(Saline)=8, n(MAM-16)=11, females: n(Saline)=8, n(MAM-16)=9],, (Figure 4C-D). Theta E-F) Graphs (top) and representative traces (bottom) showing LTP following theta-burst in HPC in females (left) and males (right). Both female (E) and male (F) MAM-treated mice exhibit significantly reduced LTP (repeated measures ANOVA, F_(1,14)_=10.1, p=.001 for females and F_(1,14)_= 8, p=.001 for males).

### Male but not female MAM-16 treated mice exhibit PFC function deficits

SZ patients and several animal models of SZ exhibit deficits in PFC function. Therefore, we investigated PFC function by examining working memory function as well as synaptic transmission and plasticity in PFC brain slices. Saline and MAM-16 treated mice were subjected to the delayed alternation task, which examines spatial working memory function. Saline and MAM-16 mice were initially trained to alternate the left and right arm in the T-maze in order to receive their food reward. No significant difference was observed in the training for the alternation procedure, for both the female and male mice (Figure 5A) (one-way ANOVA,F_(1,14)_=0.80, *p*=0.20). Once delays were introduced in the alternating procedure, female MAM-16 required the same number of trials to reach criterion compared to female saline-treated mice (Figure 5A)(one-way ANOVA, F_(1,14)_=0.20, *p*=0.30). However, this was not the case for male MAM-16 mice that required increased number of trials in order to reach criterion (one-way ANOVA, F_(1,14)_= 5.7, *p*=0.02).In addition, performance in the T-maze did not differ between female MAM-16 mice and their respective saline-treated mice (t-test, *p*=0.3), while in male MAM-16 a significant reduction was observed (t-test, *p*=0.04) compared to their respective saline-treated mice (Figure 5B).

**Figure 5:**
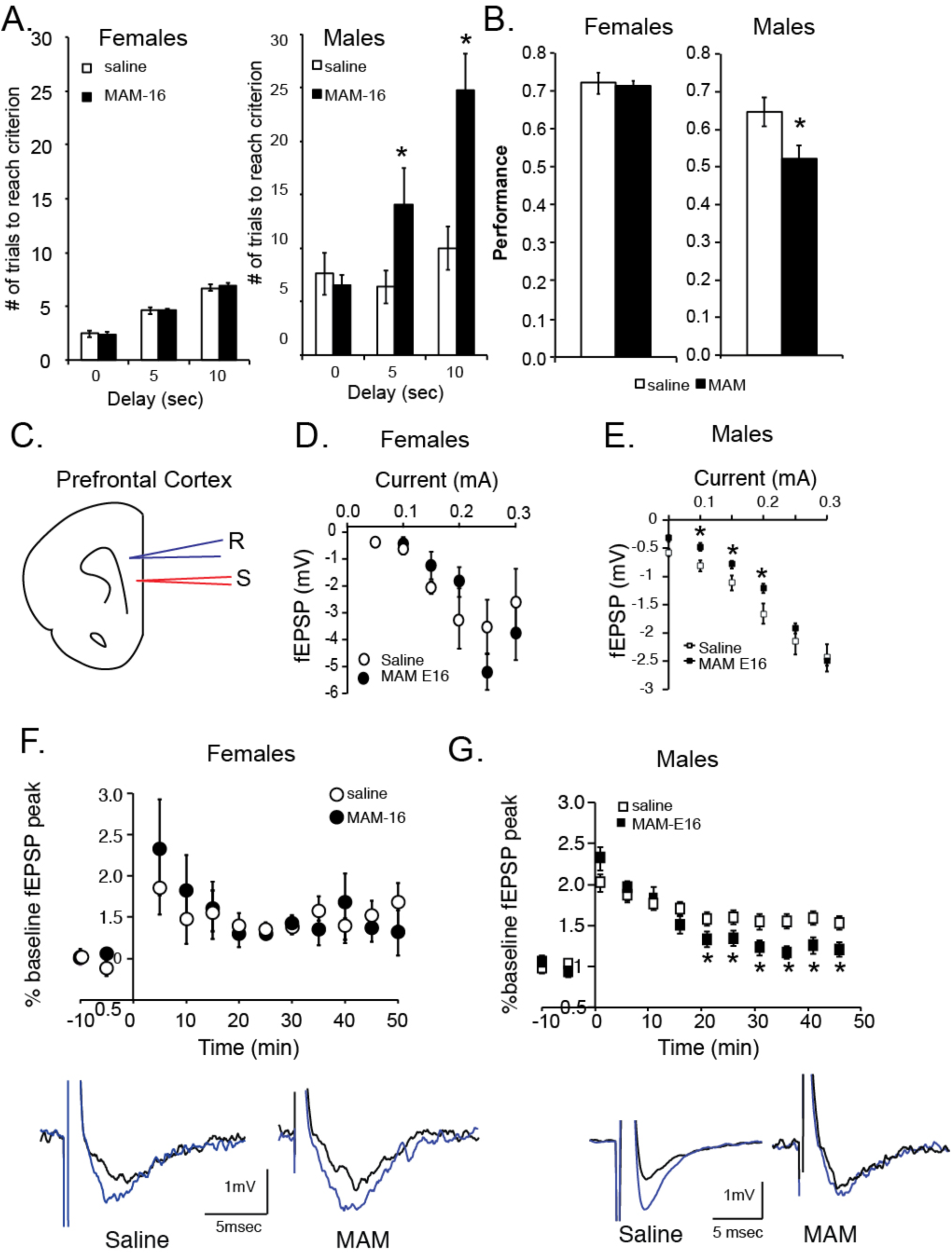
PFC function male and female MAM-16 mice. B) Graphs showing the number of cumulative trials in the delayed alternation task in the T-maze at 0, 5 and 10sec delays. No differences are present in female MAM-16 mice (n=8) compared to saline-treated (n=7) (one-way ANOVA, F_(1,13)_=0.8, p=.4), while male MAM-16 mice (n=7) require significant greater number of trials to reach criterion at the 5sec and 10sec delay B)Graphs showing that the performance (% correct trials) at the 20sec delay is significantly reduced for MAM-16 male mice (t-test, p=.02) compared to saline, but not for MAM-16 female mice (t-test, p=0.4). Illustrations indicating the position of the electrodes for the recordings in the PFC. D-E) Graphs showing the fEPSP in response to increasing current stimulation for male and female mice. No difference is found in females (repeated measures ANOVA, F_(1,14)_=0.6, p=.6) while decreased fEPSP response is observed in males (repeated measures ANOVA, ANOVA, F_(1,14)_=6.2, p=.01). F-G) Graphs (top) and representative traces (bottom) showing LTP following tetanic stimulation. There was no difference in females (repeated measure ANOVA, F_(1,14)_=0.7, p=.6), but a significant decrease is observed in males, specially from 30-50min post-tetanus (repeated measure ANOVA, F_(1,14)_=2.2, p=.0).

Working memory function is supported by both synaptic transmission and synaptic plasticity in the PFC (Miller & Cohen 2001; Blumenfeld 2006; Konstantoudaki et al. 2018). Therefore, we recorded fEPSPs from layer II/III of the PFC of saline or MAM-16 female and male treated mice (Figure 5C-D). We found no alterations in the fEPSP peak in female MAM-16 mice (Figure 4E) (repeated measures ANOVA, F_(1,14)_=0.7, *p*=0.5). However, in male MAM-16 mice the fEPSP peak was found reduced at various intensities of current stimulation (F_(1,14)_=6.2, *p*=0.01) (Figure 4D).

To determine whether synaptic plasticity in the PFC is affected, we induced LTP by tetanic stimulation. Brain slices from both saline and MAM-16 treated female mice enhanced the fEPSP following tetanic stimulation that lasted for at least 50min (repeated measures ANOVA, F_(1,14)_=0.6, *p*=0.6) (Figure 5F). On the other hand, brain slices from saline-treated male mice exhibited enhancement of the fEPSP that lasted for at least 50min, while brain slices from male MAM-16-treated mice exhibited an initial enhancement of the fEPSP, which was significantly reduced and approached baseline levels about 15 min following the tetanus (repeated measure ANOVA, F_(1,14)_=2.2, *p*=0.03)(Figure 5G). Therefore, it seems that LTP in the PFC is impaired in male MAM-16 mice but not in female MAM-16 treated mice, similar to the working memory deficits that are pronounced in male compared to female MAM-16 mice.

### Proteome analysis of the saline and MAM-treated PFC

Given the sex differences observed in the PFC of MAM-16 treated mice, we proceeded in a proteomic analysis of saline and MAM-16 treated male and female mice, in order to determine whether these sex differences can be observed at the proteome level and could explain the physiological and behavioral differences. PFC tissue was isolated from male and female saline-treated and MAM-treated mice and subjected to unbiased proteomic analyses. Venn diagrams indicate that more MAM-16 specific proteins are identified in males compared to females (Figure 6A). We next analyzed quantitative changes in three KEGG pathways affected in SZ, namely the “dopaminergic synapse”, “calcium signaling” and “glutamatergic synapse” pathways. Proteins clustered under the “dopaminergic synapse” KEGG pathway (Figure 6B) exhibit an overall tendency for down-regulation, over the respective saline-treated mice. Key proteins that are down-regulated in both males and females include tyrosine hydroxylase (TH) and monoamine oxidase B (MAOB). Proteins grouped in the “glutamatergic synapse” KEGG pathway (Figure 6C) are characterized by down-regulation that is stronger for MAM males (61% of regulated proteins in males are down-regulated and 53% in females). For example, glutamatergic AMPA receptors GRIA2 and GRIA3 are all found down-regulated in the male MAM-treated PFC, but up-regulated in females. Similarly, proteins grouped under the “calcium signaling” KEGG pathway, such as different subytpes of calcium-calmodulin kinases, are highly down-regulated over the respective saline-treated mice, mostly in male mice (Figure 6D). These observations jointly point towards a more prominent impairment in glutamate transmission in MAM-treated male PFC compared to female PFC and reinforce the male-specific synaptic plasticity and working memory deficits.

**Figure 6:**
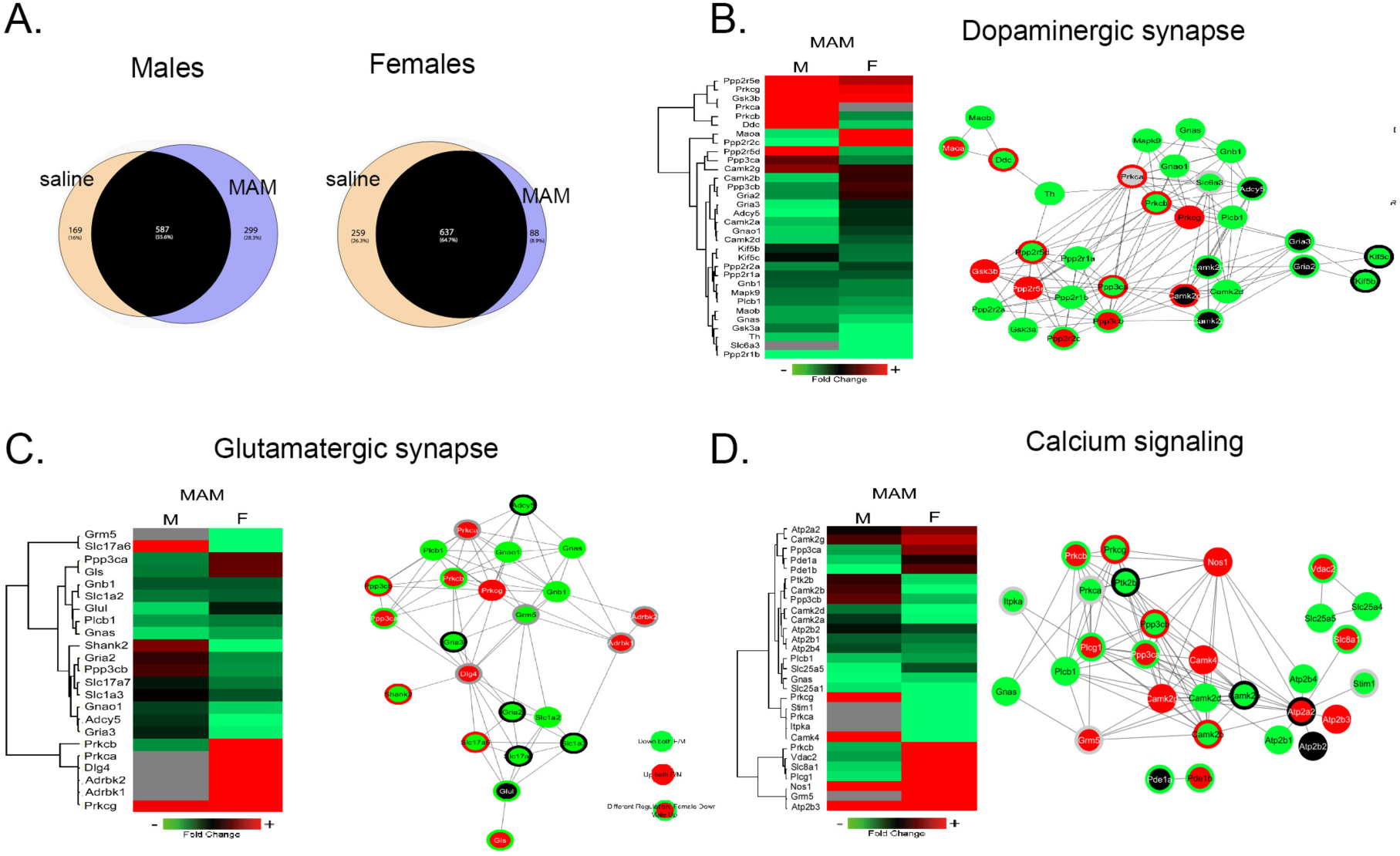
Sex differences in protein networks in PFC. A) Venn diagrams depicting the total proteome differences between the control and the MAM model in males and females. B-D) Heat-maps and protein networks of three characteristic groups of proteins found deregulated in MAM-16 female and male mice: dopaminergic synapse (B), glutamatergic synapse (C) and calcium signaling (D). The heat-maps show the Euclidean hierarchical clustering of the proteins of each group according to its fold of change in MAM-16 versus saline-treated mice, in females and males. The protein interaction network extracted from String shows the interactions (predicted and experimentally verified) for each protein in the different groups colored according to each fold of change in MAM-16 versus saline-treated groups, in females and males.

## Discussion

This study describes the SZ neurodevelopmental MAM model in mice and identifies sex differences in prefrontal cortical and hippocampal function deficits. Regarding the model, we find that the MAM model in mice is better reproduced by exposure to the mitotoxin on the 16th day of gestation, compared to the 17th day of gestation in rats. Both male and female mice exhibit thinning of the cortex and the hippocampus and indications of positive symptoms (reduced PPI in males and increased locomotor activity in response to MK-801 in females). PV reduction is only observed in male mice. HPC-dependent cognitive function is equally affected in both male and female mice, while PFC-dependent cognitive function is differentially adapted in MAM-16-treated male and female mice. The proteome of MAM-16 male PFC is differentially regulated compared to MAM-16 female PFC.

### Comparison with the MAM model in male rats

The MAM model has been established and extensively studied in male rats (Modinos et al. 2015). Our results in mice indicate that the MAM model can also be used in mice by performing the MAM treatment one-day earlier, compared to rats. Species comparison of embryonic development has indicated that the mouse embryo brain development mouse occurs faster by 1-2 days compared to the rat. Therefore, the GD 16^th^ in mice is the equivalent to GD 17^th^ of rats and corresponds to a peak proliferation stage of neural cells, which then migrate particularly in the HPC and cortex (Bayer & Altman 2004). It is expected that MAM administration after this peak stage of proliferation, would affect brain development in a lesser degree; possibly affecting only those regions that are formed later in brain development.

Our histological analysis shows thinning of the ACC in both male and female MAM-16 mice, but not MAM-17 mice. On the other hand, the HPC is affected in both MAM-16 and MAM-17 male and female mice. These findings are in agreement with the histopathological deficits found in MAM-17 rats, where both mPFC and HPC (dorsal and ventral part) have decreased thickness or total size, compared to control animals (Flagstad et al. 2004; Moore et al. 2006; Featherstone et al. 2007; Matricon et al. 2010; Hradetzky et al. 2012). Finally, no alterations in the width of BC, a sensory cortex, are found, in agreement to Moore et al.(Moore et al. 2006). However, there are reports of shrinkage in the sensory-motor cortex (Featherstone et al. 2007; Matricon et al. 2010).

Several studies have shown decreased PPI of the acoustic startle reflex in male MAM-17 rats (Moore et al. 2006; Le Pen et al. 2006; Hazane et al. 2009), which is one of the positive symptoms also observed in SZ patients. PPI is characterized as a cross species measure of sensory-motor gating and in animals can be indicative of both hyper-dopaminergic and hypo-glutamatergic function of the brain (Van Den Buuse 2010). We also find decreased PPI in MAM-16, but not in MAM-17 treated mice. Both MAM-treated groups exhibit normal responses in startle stimuli, indicating that MAM exposure has not affected either the auditory pathway or the motor output of the acoustic startle circuit; therefore the observed decrease in PPI is due to a sensorimotor gating deficit. These experiments imply a stronger effect of prenatal MAM exposure on GD16.

Another consistent finding regarding the deficiency in post-mortem studies of SZ patients is the decreased PV-immunoreactivity or mRNA levels of the PV expressing gene in the cortex, including the dorsolateral PFC, ACC, and HPC (Zhang & Reynolds 2002; Torrey et al. 2005; Konradi et al. 2011; Wang et al. 2011). Our findings showing reduced PV expression in male PFC and HPC are in agreement with human reports and with studies of MAM-17 male rats (Penschuck et al. 2006; Lodge et al. 2009). Therefore, our study proposes that the MAM-16 model in mice is comparable to the MAM-17 rat model with regards to ‘schizotypic-like’ alterations.

Both SZ patients and the MAM-17 rats exhibit significant cognitive deficits, including working memory and spatial memory deficits. Specifically, MAM-17 rats have deficits in the water-maze and Y-maze spontaneous alternation tasks (Gourevitch et al. 2004; Snyder et al. 2013; Gastambide et al. 2015; Ratajczak et al. 2015; Gill et al. 2017). In our study, male MAM-16 mice exhibit deficits in learning and performance of the delayed alternation task and in fear memory. In addition, we find significant decrease in both the fEPSP and LTP in male HPC. Studies in rats have shown reduced intrinsic excitability in MAM-17 rats, which could account for the reduced fEPSP observed in our study, but no difference in LTP (Sanderson et al. 2012). For LTP, a slightly different protocol of theta-burst was used which could account for the difference in our findings (three trains instead of 2, 10sec instead of 20sec). Furthermore, adaptations in up-down states have been observed in male MAM rats, indicative of impaired PFC function(Moore et al. 2006).

Several proteomic human studies have been conducted, mostly in blood samples, but also in cells and tissues from SZ patients, in an effort to identify biomarker candidates of SZ (for review see Nascimento & Martins-de-Souza 2015). Our proteomic analysis of PFC samples revealed alterations in two consistently affected pathways in SZ, the dopaminergic and the glutamatergic transmission pathway, as well as in the related calcium signaling pathway, but also in energy metabolism-linked proteins (glycogen synthase kinase 3b), which is also another very common dysfunction of SZ (Martins-de-Souza, Harris, et al. 2010).

### Sex differences possibly observed in humans and other animal models ofschizophrenia

Our study revealed significant sex differences in PV-expression, PFC-dependent but not HPC-dependent cognitive deficits. While the number of PV-expressing neurons decreased in male MAM-16 mice, no alterations were identified in either PFC or HPC in female mice. Since we have not measured the total number of cells in each region of interest, we cannot exclude the possibility that there is a total loss of neuronal cells in MAM-16 mice PFC and HPC, which could result in a concomitant reduction of PV-positive neurons. However, our results show that reductions in PFC and HPC occurred in both males and females, suggesting that the reduction of PV in males is a sex specific adapation. In humans, there is also indication of female patients that do not show alterations in PV-positive interneurons of HPC (Zhang & Reynolds 2002).

Although SZ affects both men and women, a higher incidence is observed in men, along with an earlier age of onset and a more severe phenotype. Particularly, men exhibit more negative and cognitive symptoms compared to females and show more severe structural brain deficits (Leung & Chue 2000; Abel et al. 2010; Ochoa et al. 2012).However, sex is not always incorporated as a variable and the vast majority of research is conducted in male subjects, leaving gaps in our knowledge regarding sex-related manifestations of the disease that could affect management.

With regards to MAM model in rats, the majority of the studies have been conducted in male rats. To our knowledge, there are three studies that evaluated female rats at the behavioral and electrophysiological level (Hazane et al. 2009; Pen et al. 2011; Snyder et al. 2013) indicating the presence of positive symptoms and reduced HPC function, in agreement to our findings. Sex differences have also been observed in other developmental animal models studies, including the Poly I:C exposure model and the neonatal ventral hippocampal lesion model (for review see Hill 2016). Our study further reports that female MAM-16 mice do not exhibit significant impairments in PFC function, as indicated by similar performance in the delayed alternation task in the T-maze, same LTP emergence and maintenance. Stronger deficits in PFC function have also been found in the Disc1 mouse model, showing enhanced excitation-to-inhibition ratio in male but not in female PFC(Holley et al. 2013). Although there is a lack of a significant number of studies that include both male and female subjects, the data so far indicate an increase in the vulnerability of the male PFC to neurodevelopmental insults (Hill 2016). Our study further shows that working memory function and synaptic transmission and plasticity are rescued in female MAM-16 mice. In addition, comparative proteomic analysis between male and female MAM-16 PFC reveals a widespread differential modulation of the proteome between the two sexes. The observed sexually dimorphic regulation of the AMPA receptor subunits A2 and A3 (GRIA2 and GRIA3) support our findings of increased vulnerability of PFC in MAM-16 male but not female mice, which is in agreement with findings from the ACC of human patients (Martins-de-Souza, Schmitt, et al. 2010). Overall, our description of the mouse MAM SZ model, from the viewpoint of sex comparisons, proves that our model mirror disease sex-specific phenotypes in human and could offer an ideal platform to study differences in pathophysiology and treatment responses to the direction of tailoring management.

### Possible mechanisms of PFC vulnerability in males but similar vulnerability of female HPC

Estrogens have been shown to exert a protective effect on the hippocampus, particularly in response to stress during adulthood. It is likely that due to the early development of the hippocampus (i.e. before adolescence), estrogens are not able to rescue HPC function in prenatal insults. On the other hand, the PFC continues to develop through adolescence at which time estrogens in females could exert their possible protective effect (Davis et al. 2016; McEwen et al. 2015). Furthermore, male sensitivity to perinatal testosterone may be interrupted by MAM exposure in a way that the brain, and particularly the PFC, is affected in adulthood (McCARTHY 2008).

The MAM mouse model having a sex differences phenotype similar to that of the human population could be used in conjuction with investigation of developmental events to shed light on different developmental changes that could be affected in males and females.

### Future directions

The validation of the MAM-16 mouse model of SZ reported in this study can contribute significantly to further testing of the two-hit hypothesis for the emergence of SZ. In support of the ‘two-hit’ hypothesis (Maynard et al. 2001; Monte et al. 2017), we propose the development of a MAM model in mice that incorporates genetic and environmental dimensions. Hence, the MAM mouse model can be used in combination with different types of transgenic mice, such as: a) mice in which specific genes or genetic polymorphisms that are associated with the emergence of SZ are modified (knocked-out or mutated), b) mice in which genetic interventions are applied conditionally at different ages in order to specify critical developmental periods for the emergence and/or prevention of SZ, c) transgenic animal models that provide tools to better delineate the pathophysiology of the disease in a neural circuit specific manner. Finally, the presence of sex differences in the MAM mouse model that resemble those of the human population allows sex-specific study design for the diagnosis and treatment of the SZ.

## Acknowledgements

This work has been funded by a NARSAD young investigator award (KS), the Special Accounts for Research of the University of Crete (KS, KC) and an ELIDEK Ph.D. scholarship (VS). We would like to thank Kiriaki Thermos for proofreading the final version of the manuscript and all the members of the Sidiropoulou lab for fruitful discussions and comments.

Author contributions
K.C. and K.S. conceived and designed research; K.C., A.V., E.K., V.S. and K.S. performed experiments; K.C., A.V., E.K., V.V., M.A., and K.S. analyzed data; K.C., E.C., and K.S. interpreted results of experiments; K.C., M.A. and K.S. prepared figures; K.C. and K.S. drafted manuscript; K.C., E.K., M.A. and K.S. edited and revised manuscript; K.C., A.V., E.K., V.S., V.V., E.C., M.A. and K.S. approved final version of manuscript.

